# COVID-19 patients upregulate toll-like receptor 4-mediated inflammatory signaling that mimics bacterial sepsis

**DOI:** 10.1101/2020.07.17.207878

**Authors:** Kyung Mok Sohn, Sung-Gwon Lee, Hyeon Ji Kim, Shinhyea Cheon, Hyeongseok Jeong, Jooyeon Lee, In Soo Kim, Prashanta Silwal, Young Jae Kim, Chungoo Park, Yeon-Sook Kim, Eun-Kyeong Jo

## Abstract

Observational studies of the ongoing coronavirus disease 2019 (COVID-19) outbreak suggest that a cytokine storm is involved in the pathogenesis of severe illness. However, the molecular mechanisms underlying the altered pathological inflammation in COVID-19 are largely unknown. We report here that toll-like receptor (TLR) 4-mediated inflammatory signaling molecules are upregulated in peripheral blood mononuclear cells (PBMCs) from COVID-19 patients, compared with healthy controls. Among the most highly increased inflammatory mediators in severe/critically ill patients, S100A9, an alarmin and TLR4 ligand, was found as a noteworthy biomarker, because it inversely correlated with the serum albumin levels. These data support a link between TLR4 signaling and pathological inflammation during COVID-19 and contribute to develop therapeutic approaches through targeting TLR4-mediated inflammation.

## Introduction

The coronavirus disease 2019 (COVID-19) is caused by the novel coronavirus SARS-CoV-2 and has spread globally causing international concerns (*1-3*). As of July 16, 2020, there were 13,378,853 confirmed cases of COVID-19 in 216 countries, with 580,045 confirmed deaths (*4*). Most patients are asymptomatic or recover after mounting a self-limiting antiviral response with the development of neutralizing anti-viral antibodies and cell-mediated immunity (*5*). However, ∼10% of all cases become serious, with dyspnoea, lymphopenia, and extensive chest x-ray abnormalities and half of these become critically ill, with respiratory and multi-organ failure (*6-8*). There appears to be a relationship between the clinical and immunological features of COVID-19, as the disease severity correlates with certain immunological markers (*7, 9*). Recent studies have shown that severe/critically ill (SEVERE) patients exhibit “cytokine storm”, which is related to the production of excessive cytokines, dysregulated immune cell function, and massive systemic inflammation (*1, 5, 10*). Understanding the causes of altered immune features of COVID-19 would enable the refinement of preventive vaccine targets and accelerate therapeutic development. Despite this, the molecular mechanisms underlying exaggerated inflammatory phenotypes during COVID-19 are largely unknown. We herein show that toll-like receptor (TLR) 4-mediated inflammatory signaling molecules, which mimic pathogenesis of bacterial sepsis, are upregulated in peripheral blood mononuclear cells (PBMCs) from COVID-19 patients.

## Results and discussion

### Characterization of immune features of COVID-19 patients in terms of clinical severity

To investigate the immune signaling signature of COVID-19, a total of 48 Korean subjects [untreated COVID-19 patients with various clinical severities (n = 28) and healthy controls (n = 20)] were enrolled in the study. Table 1 summarizes the characteristics and laboratory findings of 20 mild/moderate [MILD; median age 53.5 (range 21–97) years] and eight SEVERE [63.5 (range 36–78) years] patients. This study was approved by Institutional Research and Ethics Committee at Chungnam National University Hospital (CNUH 2020-03-056; Daejeon, Korea) and was conducted in accordance with the Declaration of Helsinki (*11*). In the SEVERE group, seven of eight (87.5%) patients had fever at the time of sampling versus only two (10%) in the MILD group. The median time from symptom onset to mechanical ventilation was 10 (range 8–12) days. Underlying comorbidities were present in about half of the patients (hypertension, diabetes mellitus, dementia, schizophrenia, and chronic kidney disease) and did not differ between the groups. The median time from symptom onset to sampling was 5.5 (range 5–10) and 5.7 (range 5– 11) days in the MILD and SEVERE groups, respectively. The scores for the degree of illness were higher in the SEVERE group at the time of sampling. The Charlson comorbidity index was similar in the two groups, because age was matched and the underlying conditions did not differ significantly. All MILD patients recovered fully without sequelae, while four SEVERE patients required extracorporeal membrane oxygenation. One patient died of persistent pneumonia and septic shock. Among the laboratory parameters, hypoalbuminemia, a high neutrophil-to-lymphocyte ratio, and increased serum C-reactive protein and lactate dehydrogenase levels were associated with disease severity (Table 1).

**Table 1.**
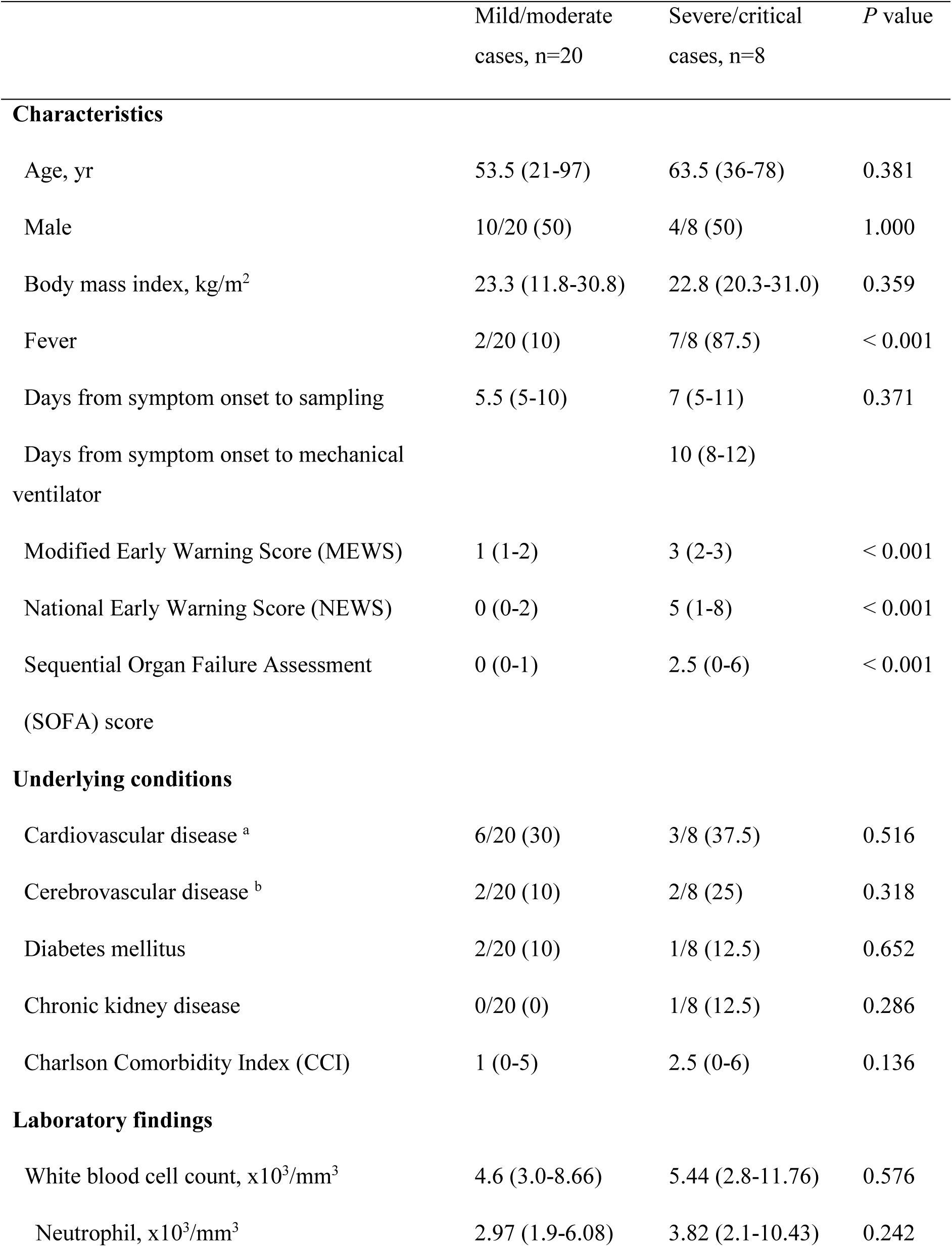
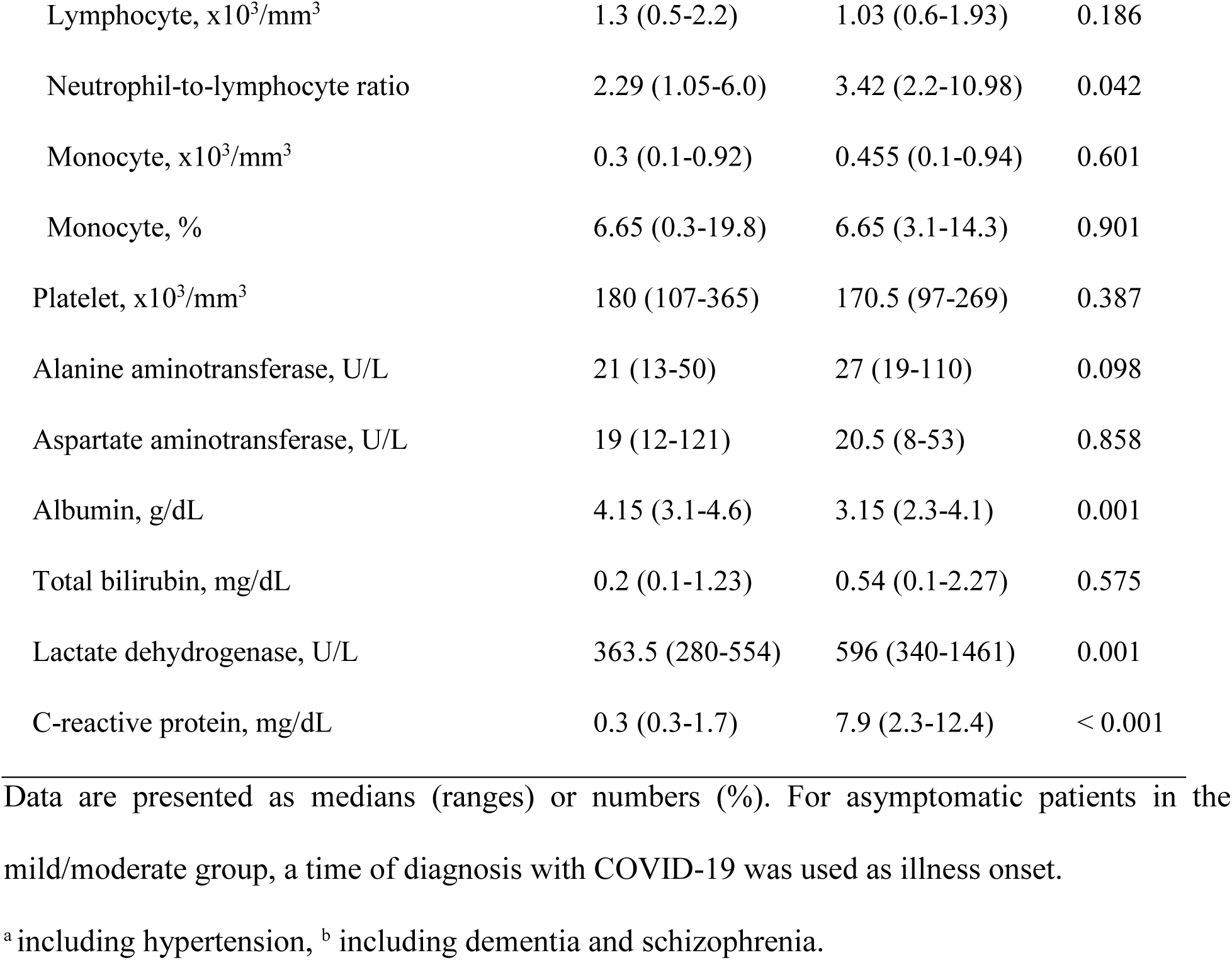
Characteristics and laboratory findings of patients with COVID-19.

|  | Mild/moderate<br>cases, n=20 | Severe/critical<br>cases, n=8 | *P* value |
| --- | --- | --- | --- |
| **Characteristics** |  |  |  |
| Age, yr | 53.5 (21-97) | 63.5 (36-78) | 0.381 |
| Male | 10/20 (50) | 4/8 (50) | 1.000 |
| Body mass index, kg/m<sup>2</sup> | 23.3 (11.8-30.8) | 22.8 (20.3-31.0) | 0.359 |
| Fever | 2/20 (10) | 7/8 (87.5) | < 0.001 |
| Days from symptom onset to sampling | 5.5 (5-10) | 7 (5-11) | 0.371 |
| Days from symptom onset to mechanical ventilator |  | 10 (8-12) |  |
| Modified Early Warning Score (MEWS) | 1 (1-2) | 3 (2-3) | < 0.001 |
| National Early Warning Score (NEWS) | 0 (0-2) | 5 (1-8) | < 0.001 |
| Sequential Organ Failure Assessment (SOFA) score | 0 (0-1) | 2.5 (0-6) | < 0.001 |
| **Underlying conditions** |  |  |  |
| Cardiovascular disease <sup>a</sup> | 6/20 (30) | 3/8 (37.5) | 0.516 |
| Cerebrovascular disease <sup>b</sup> | 2/20 (10) | 2/8 (25) | 0.318 |
| Diabetes mellitus | 2/20 (10) | 1/8 (12.5) | 0.652 |
| Chronic kidney disease | 0/20 (0) | 1/8 (12.5) | 0.286 |
| Charlson Comorbidity Index (CCI) | 1 (0-5) | 2.5 (0-6) | 0.136 |
| **Laboratory findings** |  |  |  |
| White blood cell count, x10<sup>3</sup>/mm<sup>3</sup> | 4.6 (3.0-8.66) | 5.44 (2.8-11.76) | 0.576 |
| Neutrophil, x10<sup>3</sup>/mm<sup>3</sup> | 2.97 (1.9-6.08) | 3.82 (2.1-10.43) | 0.242 |
| Lymphocyte, x10<sup>3</sup>/mm<sup>3</sup> | 1.3 (0.5-2.2) | 1.03 (0.6-1.93) | 0.186 |
| Neutrophil-to-lymphocyte ratio | 2.29 (1.05-6.0) | 3.42 (2.2-10.98) | 0.042 |
| Monocyte, x10<sup>3</sup>/mm<sup>3</sup> | 0.3 (0.1-0.92) | 0.455 (0.1-0.94) | 0.601 |
| Monocyte, % | 6.65 (0.3-19.8) | 6.65 (3.1-14.3) | 0.901 |
| Platelet, x10<sup>3</sup>/mm<sup>3</sup> | 180 (107-365) | 170.5 (97-269) | 0.387 |
| Alanine aminotransferase, U/L | 21 (13-50) | 27 (19-110) | 0.098 |
| Aspartate aminotransferase, U/L | 19 (12-121) | 20.5 (8-53) | 0.858 |
| Albumin, g/dL | 4.15 (3.1-4.6) | 3.15 (2.3-4.1) | 0.001 |
| Total bilirubin, mg/dL | 0.2 (0.1-1.23) | 0.54 (0.1-2.27) | 0.575 |
| Lactate dehydrogenase, U/L | 363.5 (280-554) | 596 (340-1461) | 0.001 |
| C-reactive protein, mg/dL | 0.3 (0.3-1.7) | 7.9 (2.3-12.4) | < 0.001 |

Data are presented as medians (ranges) or numbers (%). For asymptomatic patients in the mild/moderate group, a time of diagnosis with COVID-19 was used as illness onset.
<sup>a</sup> including hypertension, <sup>b</sup> including dementia and schizophrenia.

In this study, we first assessed the profiles of immunological determinants depending on disease severity in Korean COVID-19. To examine the immune-related transcriptome profiles induced by COVID-19 infection, we performed nCounter Human Immunology gene expression assays for PBMCs from eight SEVERE and 20 MILD patients and 20 healthy controls (HC) (Fig. 1A). Using principal component analysis (PCA), we found that HC was clearly separated from both SEVERE and MILD, while the two patient groups intermingled (Fig. 1B). These data imply that altered expression of immune-related genes is a transcriptional hallmark of COVID-19 and that the overall immune transcriptome profiles are similar in SEVERE and MILD groups in Korea. To determine which genes are differentially expressed in COVID-19 patients, we compared the expression of immune-related genes between HC and COVID-19 patients. In all, 298 differentially expressed immune genes (DEiG) were identified, and they are mainly involved in cytokine– cytokine receptor interaction and nuclear factor (NF)-κB signaling pathways (Fig. 1C, D). The same analysis was repeated for each patient group separately, and we identified 230 and 255 DEiGs in the SEVERE and MILD samples, respectively (Fig. 1A, C). The cytokine–cytokine receptor interaction was the top enriched pathway in both the SEVERE and MILD groups (Fig. S1B, D).

**Fig. 1.**
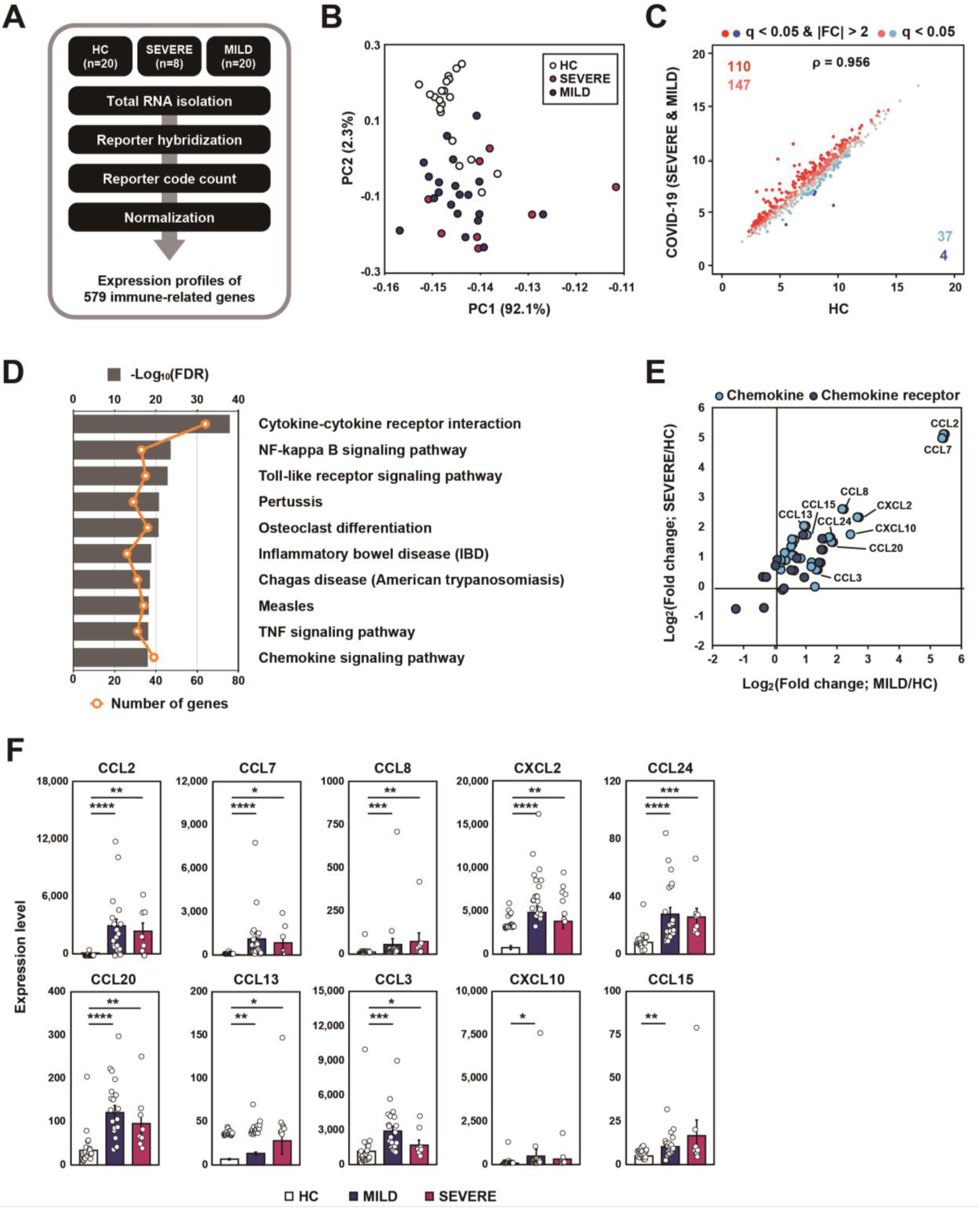
Transcriptome analysis reveals that immune gene expression profiles of COVID-19 patients are distinct to HC. (A) Schematic diagram of the immune transcriptome analysis in this study. (B) A result of principal component analysis of log_2_-transformed 579 immune gene expression levels. (C) The scatter plots representing 579 immune genes with the log2-transformed FPKM for COVID-19 patients compared to HC. (D) The ten most significantly enriched KEGG pathways of the 298 DEiGs from COVID-19 patients compared to HC. (E) Log_2_-transformed fold changes of chemokine and chemokine receptor genes from MILD (x-axis) and SEVERE (y-axis) versus HC. (F) Expression levels (FPKM) of marked chemokines in (E). Error bars indicates S.E.M. *P < 0.05, **P < 0.01, ***P < 0.001, ****P < 0.0001. *P* values were calculated using Mann-Whitney U test and adjusted P values (FDR) were shown.

We then investigated the full list of gene families associated with the cytokine–cytokine receptor interaction pathway. Using the HUGO gene nomenclature database, members of several gene families were identified, including chemokines, interleukin (IL), tumor necrosis factor (TNF), and interferon (Fig. S2A). Notably, C-C motif (CC) chemokines [CC chemokine ligand (CCL) 2, CCL7, CCL8, CCL24, CCL20, CCL13, and CCL3], C-X-C motif (CXC) chemokines [CXC chemokine ligand (CXCL) 2 and CXCL10], and chemokine receptor subfamilies, were most numerous, and were significantly (FDR < 0.05) upregulated in both MILD and SEVERE COVID-19 patient groups (Fig. 1E, F; Fig. S2B). These data partially correlated with recent reports that severe and critically ill cases are associated with defective immune responses, i.e. lymphopenia, high neutrophil-to-lymphocyte ratio, and increased inflammatory cytokine and chemokine levels (*1, 5, 12-17*). However, our results are unique in that we found that many CC chemokines and ILs are highly upregulated in both MILD and SEVERE patients, compared with HC. Together, these data suggest that abnormal inflammatory chemokine generation represents common immune features during COVID-19.

### Elucidation of molecular signaling pathways of immune transcriptome during COVID-19

TLR signaling pathways are mediated by the components, including sensors that recognize certain pathogen- and damage-associated molecular patterns (PAMPs and DAMPs) and the adaptors that transduce signals (*18*). Among TLRs, TLR4 can recognize lipopolysaccharide (LPS), other PAMPs, and DAMPs, at the cell surface, whereas TLR3, TLR7, TLR8, and TLR9 are exclusively expressed in endosomal compartments and recognize viral components (*18-20*). TLR4 is the only TLR to transduce innate immune signals through both myeloid differentiation primary-response 88 (MyD88) and Toll-IL-1 receptor-domain-containing adaptor-inducing IFN-β (TRIF) to activate NF-κB and interferon regulatory factor (IRF) signaling, respectively (*18-20*).

Next, we explored the second top hit pathway, NF-κB signaling (Fig. 1D), which is one of the major hyper-activated pathways following COVID-19 infection (*21*). With some exceptions (such as LCK, CD40LG, PLAU, PTGS2, and TRAF5), the expression of TLR4 and its related/downstream signaling molecules (CD14, MYD88, IRAK1, TRAF6, TIRAP, TICAM) were significantly (FDR < 0.05) upregulated (Fig. 2A). In addition, most NF-κB signaling pathway genes (NFKBIA, NFKB1, RELA, NFKB2) were significantly (FDR < 0.05) upregulated (Fig. 2A). These data suggest that TLR4-mediated NF-κB signaling pathway activation is involved in the upregulation of inflammatory responses in patients with COVID-19 infection. Interestingly, there were no significant differences in the expression of IRF3, TLR3, TLR7, TLR8, and TLR9, all of which are related to putative viral signaling (*22, 23*), between COVID-19 patients and HC (Fig. 2B). These data strongly suggest that, TLR4-triggered inflammatory signaling, rather than viral stimulation, contribute to molecular pathogenesis during COVID-19. We also found that IL-1β and its downstream inflammatory signaling molecules (IL1R1, MYD88, IRAK1, TRAF6, NFKBIA, NFKB1, RELA) were dramatically elevated in COVID-19 patients (Fig. 2A). NF-κB signaling pathway is required for proinflammatory cytokine/chemokine generation and the production of antimicrobial proteins (*20, 24*); IL-1 family members are involved in the initiation of potent inflammatory responses, orchestration of innate and adaptive immunity, and development of sepsis (*25, 26*). Although both TLR and IL-1 signaling activation is critical for innate immune defense against a variety of pathogens, dysregulation of this signaling pathway can lead to pathogenesis of various diseases including inflammatory and autoimmune diseases (*25-28*). The data suggest that the upregulated profiles of TLR4, IL1R, and NF-κB signaling pathway molecules in COVID-19 patients are presumably associated with the altered immune responses to viral components, host DAMP signals, or cytokine signaling activation (*10*), and may contribute to uncontrolled pathological inflammation.

**Fig. 2.**
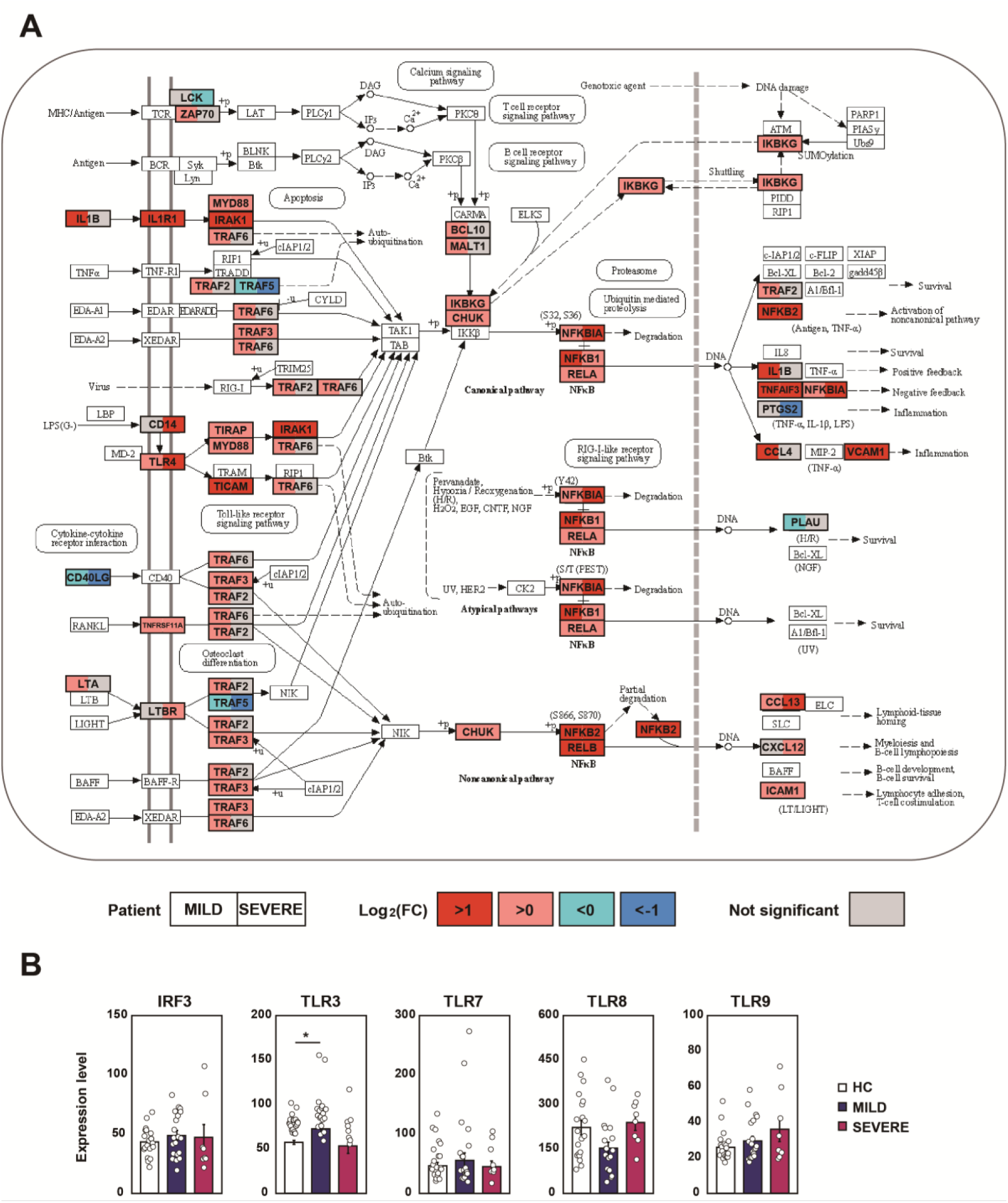
COVID-19 infection boosts NF-κB signaling pathway. **(A)** The left (MILD) and right (SEVERE) sides of box represent the mean fold change in mRNA levels, compared with HC. The NF-κB signaling pathway was adopted from KEGG database (accession number: hsa04064). **(B)** The expression levels of IRF3, TLR3, TLR7, TLR8 and TLR9 were represented by FPKM. Error bar indicates S.E.M. \**P* < 0.05. *P* values were calculated using Mann-Whitney U test and adjusted *P* values (FDR) were shown.

### Identification of SEVERE-specific immune target genes

To understand the pathophysiological differences between SEVERE and MILD patients better, we then examined whether there are any transcriptomic differences between the two COVID-19 patient groups. Directly comparing the gene expression profiles between the SEVERE and MILD groups, no genes were significantly (FDR < 0.05) differentially expressed, which might have been partly hindered by the heterogeneity of the presentation of disease severity. Therefore, we performed two pairwise transcriptome comparisons: SEVERE versus HC and MILD versus HC. From these comparisons, 58 DEiGs were identified showing significant (FDR < 0.05) changes in gene expression between SEVERE and HC, which are potential therapeutic targets, but none between MILD and HC (Fig. 3A). Intriguingly, the highly expressed SEVERE-specific upregulated genes were mainly associated with complement activation (C9 and C1QA), autoimmunity (AIRE and PRKCD), and inflammatory processes (CXCL11, CCL16, and S100A9) (Fig. 3B). The SEVERE-specific downregulated genes were linked to major histocompatibility complex proteins (HLA-DRA and HLA-DMA), T-cell factor/lymphoid enhancer-binding factor family (TCF-7 and LEF1), and natural killer cell functions (KLRB1, KLRG1, CD160, and GZMK) (Fig. 3C).

**Fig. 3.**
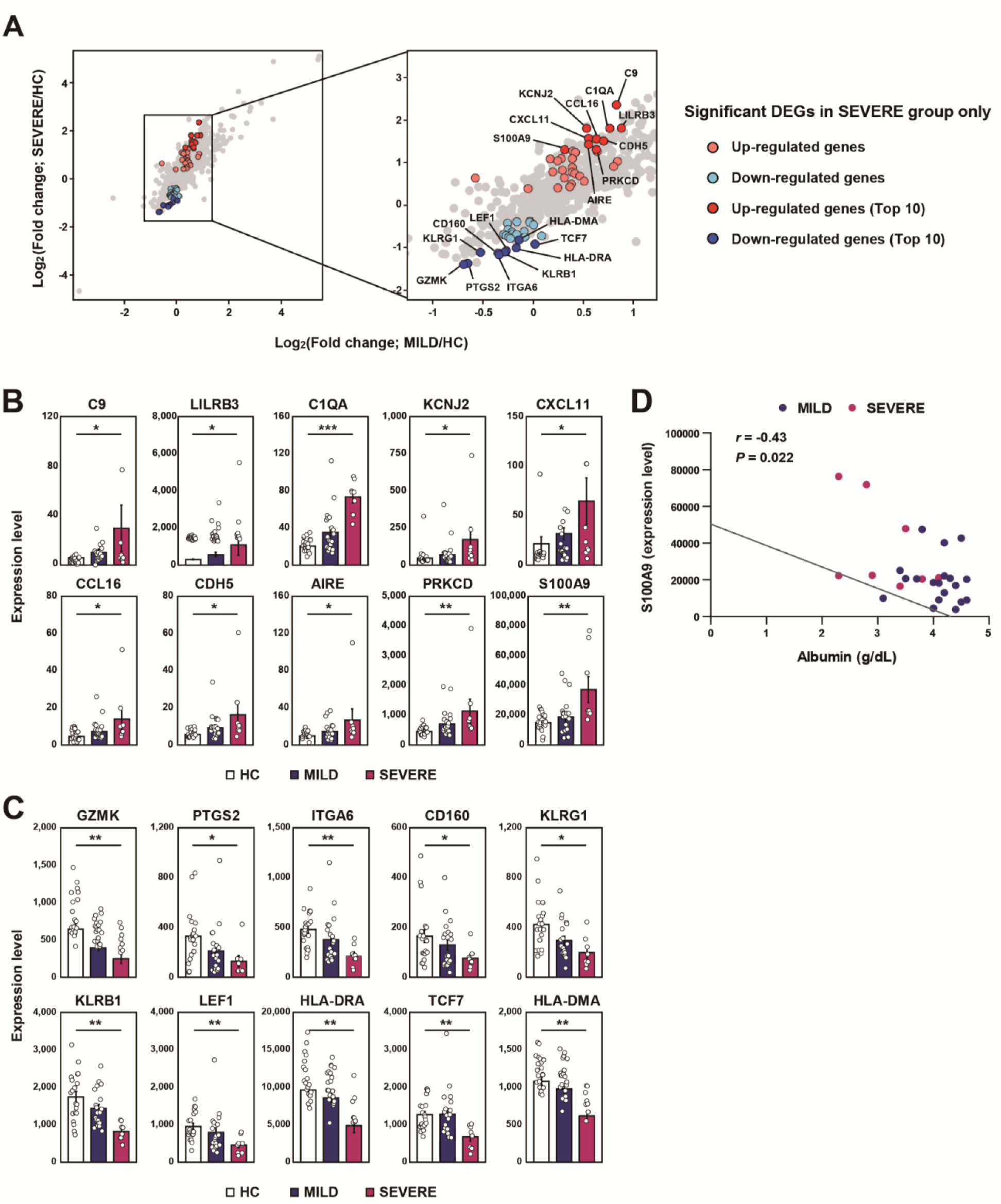
Top 10 most significantly up- and down-regulated DEiGs in SEVERE patients. **(A)** Log2-transformed fold changes of 579 immune-genes from MILD versus HC (x-axis) and SEVERE versus HC (y-axis). The genes for red and blue colors indicate up- and down-regulated DEiGs in SEVERE patients, respectively. **(B**,**C)** Comparisons of expression levels of top 10 up- (B) and down- (C) regulated DEiGs. The expression level was represented by FPKM. Error bars indicates S.E.M. **(D)** Correlation analysis between S100A9 expression level and serum albumin in MILD and SEVERE patients. Error bar indicates S.E.M. \**P* < 0.05, \*\**P* < 0.01, \*\*\**P* < 0.001, \*\*\*\**P* < 0.0001. *P* values were calculated using Mann-Whitney U test and adjusted *P* values (FDR) were shown (B, C) and Spearman’s correlation is shown (D).

We then evaluated the correlation between the immune mediators and clinical parameters (Fig. S3) and among the immune markers (Fig. S4) in COVID-19 patients. On examining the relationship between immune markers and clinical parameters (Fig. S3), we identified S100A9 as an important biomarker that was inversely correlated with serum albumin in the SEVERE group (Fig. 3D). The increased S100A9 seems to be critically important because hypoalbuminemia is associated with disease severity in COVID-19 patients (*7*). S100A8/A9 (a heterodimer complex of S100A8 and S100A9 proteins) (*29*) is a DAMP signal as a TLR4 ligand (*30*). The elevated expression of S100A8/A9 is induced by inflammation, and secreted S100A8/A9 further amplifies inflammatory soluble cytokines/chemokines, forming a feed-forward loop affecting the persistent inflammation (*31*). Importantly, S100A8/A9, as a ‘soil signal’, mediates metastasis of melanoma or breast cancers to the lung (*30*). Since S100A8/A9 protein is involved in the pathogenesis of numerous inflammation-associated and autoimmune diseases (*30, 32*), our findings provide new insight into the pathogenesis of COVID-19, and may contribute to therapeutic approaches based on the S100A9-CC chemokine-mediated inflammatory signaling. Recent promising results suggest that dexamethasone has beneficial effects to reduce deaths of patients receiving invasive ventilation or oxygen (*33*). Our findings provide a rationale to use dexamethasone, a ligand for glucocorticoid receptor, which interferes with TLR-dependent inflammatory signaling through multiple mechanisms (*34, 35*). Collectively, these data will contribute to better management and the development of targeted approaches for COVID-19 patients by enhancing our knowledge on molecular pathogenesis of the disease.

## Materials and Methods

### Study population

COVID-19 patients were confirmed by real-time quantitative polymerase chain reaction (RT-qPCR) for SARS-CoV-2 in nasopharyngeal and oropharyngeal swab, with or without sputum. Patients were categorized into two groups; mild/moderate versus severe/critical cases. In severity assessment, the World Health Organization’s COVID-19 disease severity definition was used (*36*). Twenty-eight COVID-19 patients (8 SEVERE versus 20 MILD) admitted to Chungnam National University Hospital, and age/sex-matched 20 healthy controls, giving specific informed consent were included in the study. We excluded patients with age under 19. In the severe/critical group, two patients were transferred from a long term care facility in which had a COVID-19 outbreak. They had been hospitalized with well-controlled schizophrenia. Another patient was referred to our hospital in a state of endotracheal intubation. The patients’ characteristics, clinical symptoms and laboratory test results are summarized in Table 1. All clinical and laboratory parameters were those at the time of sampling. The sampling point (median 5-6 days after illness onset) was determined by previous reports about COVID-19 patients (*3, 17*), which is relatively early in the clinical courses. In asymptomatic patients, a screened date for COVID-19 because of strong epidemiologic link was used for illness onset.

### Ethics statement

This study was approved by the Institutional Research and Ethics Committee at Chungnam National University Hospital (CNUH 2020-03-056; Daejeon, Korea) and conducted in accordance with the Declaration of Helsinki.

### Nanostring nCounter assay

Nanostring nCounter Human Immunology gene expression assays and Human miRNA expression assays were performed at PhileKorea Technology (Daejeon, South Korea), using the NanoString nCounter GX Human Immunology Kit V2 (NanoString Technologies, Inc., Seattle, WA, USA). Normalization of gene expression levels was performed by scaling with the geometric mean of the built-in control gene probes for each sample. Differentially expressed immune genes (DEiGs) among HC, SEVERE and MILD patients, were satisfied false discovery rate (FDR) < 0.05 which is analyzed and corrected by wilcox.test and p.adjust functions, respectively, implemented in stat package of R (v. 3.6.2).

### Cell Culture and RNA extraction

Human PBMCs from healthy volunteers were isolated from heparinized venous blood using Ficoll-Hypaque (Lymphoprep; Alere technologies, Oslo, Norway) as described previously (*37*). Total RNA from PBMCs was extracted using QIAzol lysis reagent (Qiagen, Hilden, Germany) and miRNeasy Mini Kits (Qiagen) according to the manufacturer’s instructions, followed by RNA quantitation.

### Bioinformatics analysis

The Spearman’s correlation coefficients of gene expressions levels of 579 immune genes were calculated with a cor.test function implemented in stat package of R. The KEGG pathway enrichment analysis was performed using DAVID (version 6.8, https://david.ncifcrf.gov) with a human reference gene set. We picked out significantly enriched pathways with FDR < 0.05.

To identify chemokine, interleukin, TNF, interferon and those receptor gene families, we downloaded gene family annotations from HUGO Gene Nomenclature Committee (https://www.genenames.org) (*38*).

## Statistical analysis

Statistical analyses were performed with Analyse-it, version 5.1 (Analyse-it Software, Ltd., Leeds, UK), SPSS Statistics for Windows, version 24.0 (SPSS Inc., Chicago, IL, USA), and GraphPad Prism, version 5.0 (GraphPad Software, San Diego, CA, USA). The data were processed by principal component analysis, Spearman’s correlation, Student’s t-test, Mann-Whitney U test, ANOVA, and Kruskal-Wallis H test, as appropriate, and detailed in each figure and figure legends. Results are presented as medians (ranges) or means ± S.E.M. (standard error of the mean) or ± S.D. (standard deviation) as indicated in figure legends. *P* < 0.05 was considered significant and marked as \**P* < 0.05, \*\**P* < 0.01, \*\*\**P* < 0.001, \*\*\*\**P* < 0.0001.

## Acknowledgments

We thank Dr. H.W. Suh and S.M. Jeon for excellent technical assistance. This work is supported by the National Research Foundation of Korea (NRF) Grant funded by the Korean Government (MSIP) (2017R1A5A2015385).

## Author Contributions

Y.S.K., K.M.S., S.G.L., H.J.K., C.P., E.K.J. participated in the research design, data curation, statistical analysis, or paper writing; H.J.K., Y.J.K., and P.S. performed experiments and analysis; Y.S.K., K.M.S., S.C., H.J., and J. L. collected blood samples from consented subjects and clinical information; I.S.K., S.G.L., C.P. did software analysis; Y.S.K., C.P., and E.K.J. supervised the study or funding acquisition. All authors read and approved the final manuscript.

## Competing interests

The authors have declared that no conflict of interest exists.

## Supplementary Figures and Figure legends

**Fig. S1.**
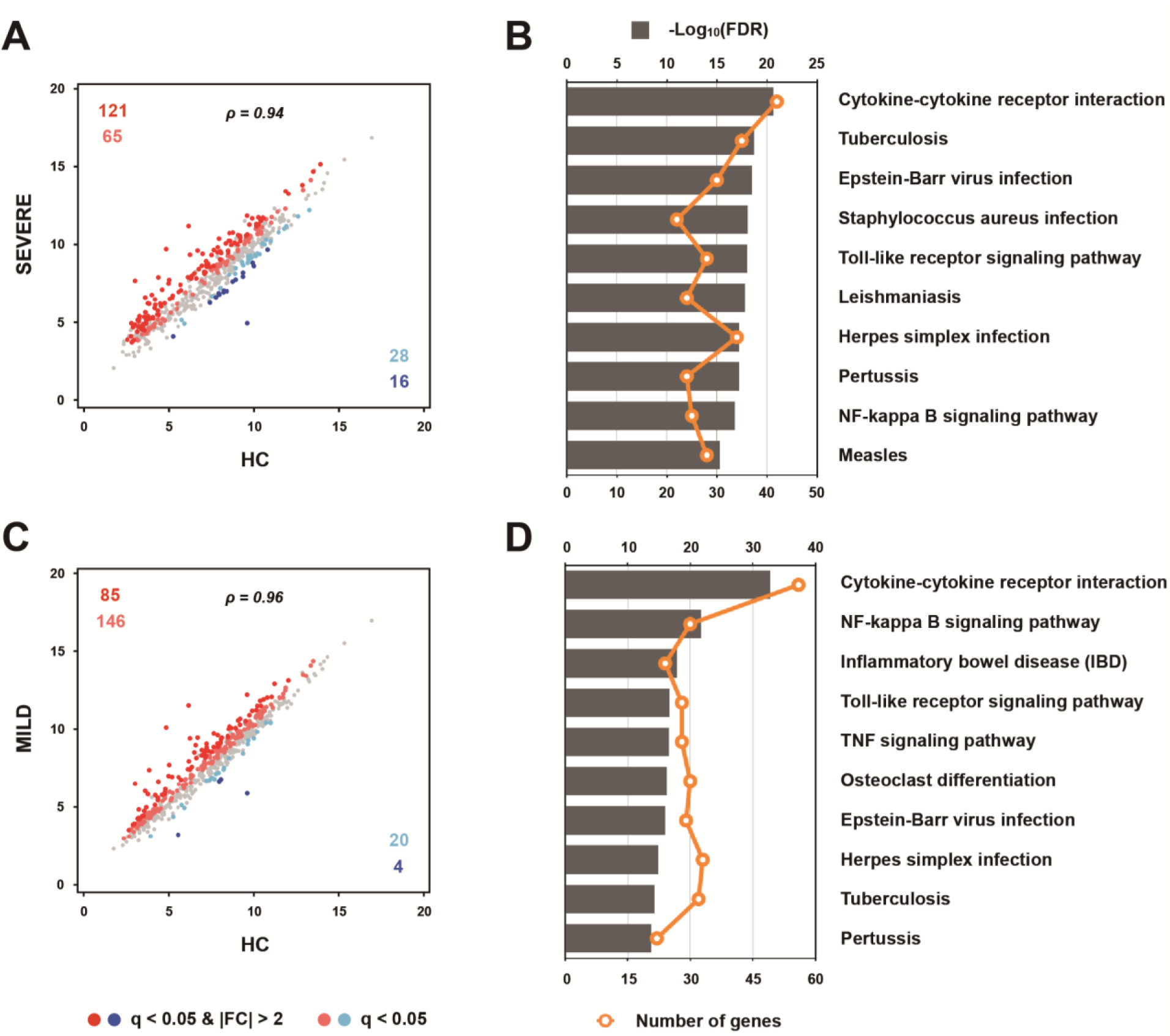
Immune gene expression profile in SEVERE and MILD COVID-19 patients. **(A**,**C)** Scatter plots representing 579 immune genes with the log2-transformed FPKM for (A) SEVERE and (C) MILD patients compared to HC. (**B**,**D)** Top 10 significantly enriched KEGG pathways associated with 230 and 255 DEiGs between (B) SEVERE versus HC and (D) MILD versus HC, respectively. *P* values were calculated using Mann-Whitney U test and adjusted *P* values (FDR) were shown. \**P* < 0.05, \*\**P* < 0.01, \*\*\**P* < 0.001, \*\*\*\**P* < 0.0001.

**Fig. S2.**
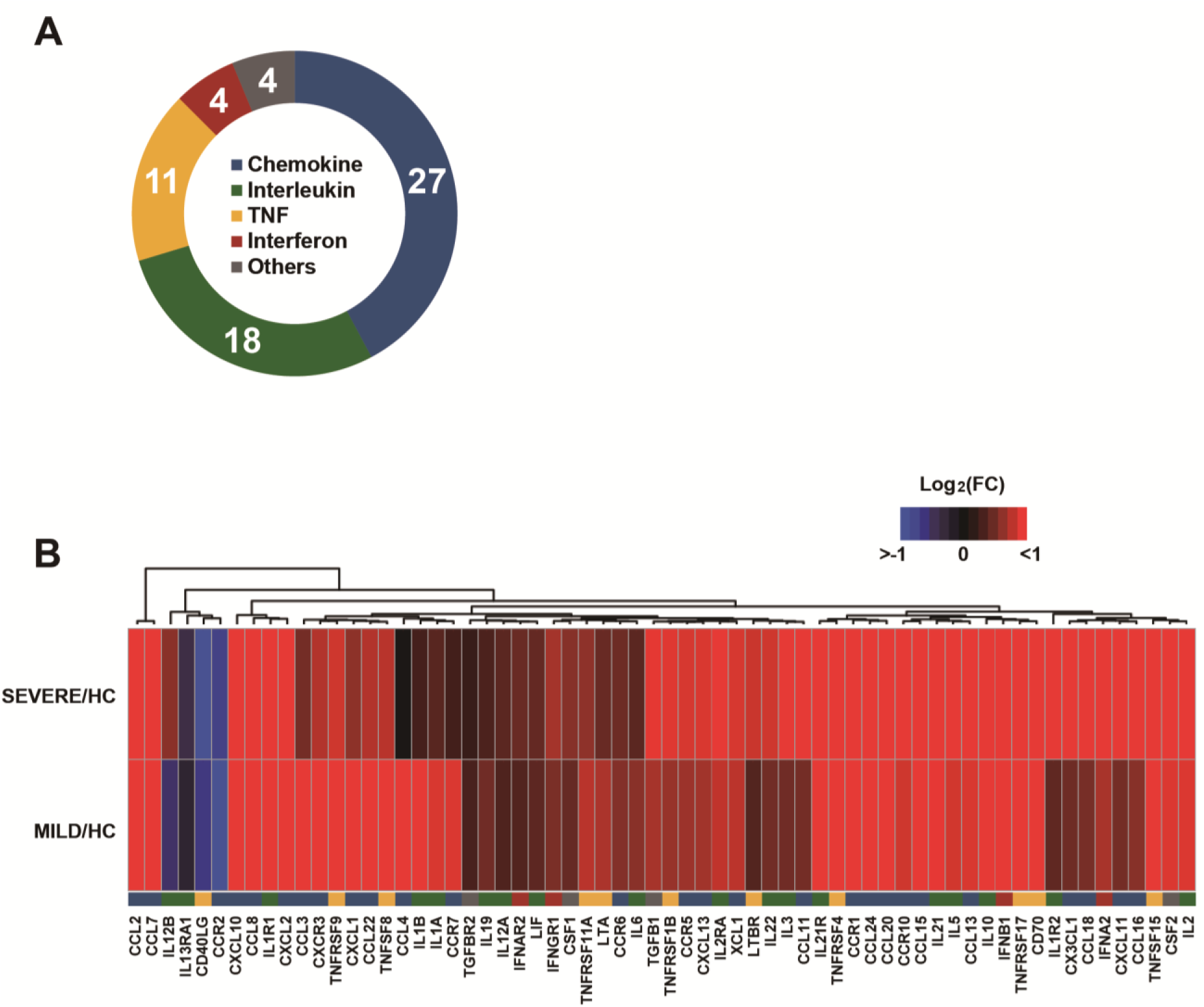
Gene expression profile of cytokine-cytokine receptor interaction pathway. **(A)** Annotation of gene families involved in the cytokine-cytokine receptor interaction pathway. (**B)** Heatmap representing log2-transformed fold changes of 64 DeiGs belonging to the cytokine-cytokine receptor interaction pathway. The hierarchical clustering was performed with Euclidean distance matrix by the hclust function implemented in stat package in R. *P* values were calculated using Mann-Whitney U test and adjusted *P* values (FDR) were shown.

**Fig. S3.**
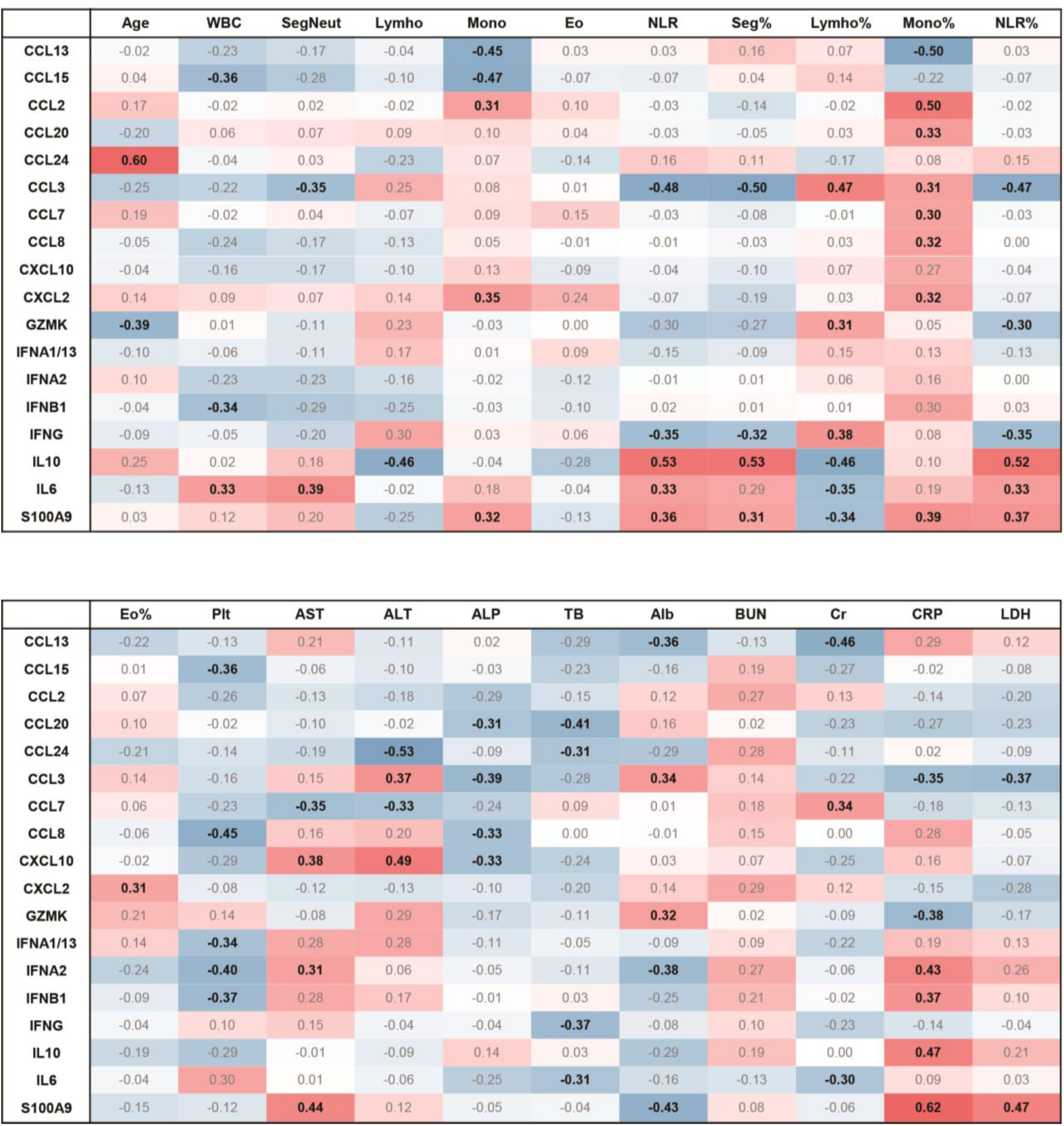
Correlation matrix between immune mediators and clinical parameters in COVID-19 patients. WBC=white blood cells, SegNeut=segmented neutrophils, Lympho=lymphocytes, Mono=monocytes, Eo=eosinophils, NLR=neutrophil-lymphocytes ratio, Plt=platelets, AST= aspartate aminotransferase, ALT =alanine aminotransferase, ALP=alkaline phosphatase, TB=total bilirubin, Alb=albumin, BUN=blood urea nitrogen, Cr=creatinine, CRP=c reactive protein, LDH=lactate dehydrogenase

**Fig. S4.**
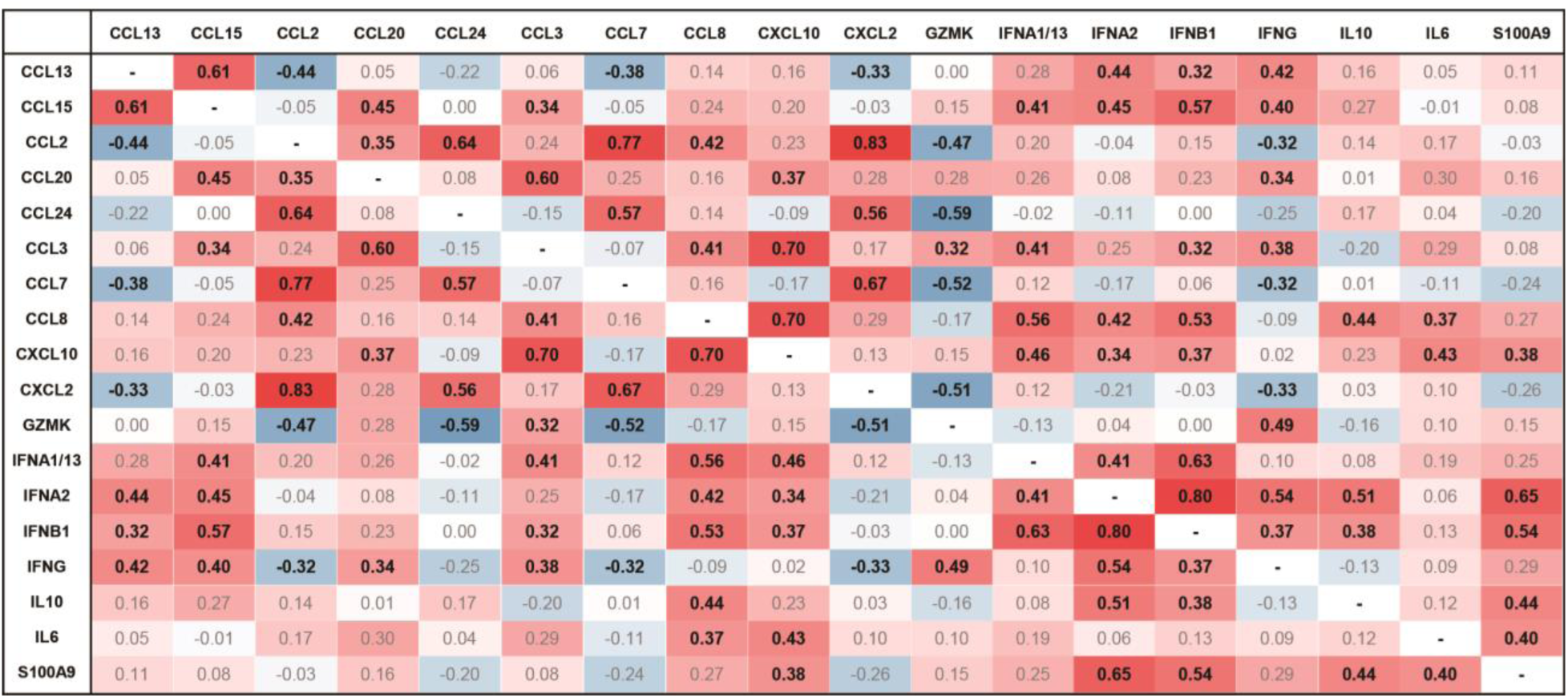
Correlation matrix among the immune parameters in COVID-19 patients.

## Notes

### Competing Interest Statement

The authors have declared no competing interest.

